# A pair of NLRs coordinately modulates NRC3-mediated ETI responses to facilitate age-dependent immunity

**DOI:** 10.1101/2022.12.20.521344

**Authors:** Xiaohua Dong, Xiaoyan Zhang, Zhiyuan Yin, Jialu Li, Chuyan Xia, Weiye Pan, Yaning Zhao, Maofeng Jing, Jinding Liu, Gan Ai, Daolong Dou

## Abstract

Two NLRs (Nucleotide-binding and Leucine-rich repeat Receptors) adjacent to each other on a locus, termed as paired NLRs, may act separately for effector recognition and subsequent signaling activation to mediate effector-triggered immunity (ETI) in many plants. However, it is largely unknown about their distribution and functions in Solanaceae species, in which NLR-Hs (Helpers NLR REQUIRED FOR CELL DEATHs) have been extensively studied. Here, we identified paired NLRs in Solanaceae species and found they harbor paired NLRs ranging from 6 to 100, which are significantly negatively correlated with the numbers of NLR-Hs. *N. benthamiana* has six paired NLRs, among which silencing of *NRCX* exhibits phenotypes of dwarfism and accelerated senescence. Importantly, *NRCX*-silencing phenotypes could be restored by simultaneously silencing its head-to-head NLR pair, thus we named it as *NRCY*. NRCX/Y pair is specific in Solanaceae species. NRCY contains non-canonical walker B and MHD motifs, but could not induce autoactive cell death in *N. benthamiana*. Instead of that, silencing *NRCY* impaired cell death triggered by Sw5b-Nsm and NRC3^D480V^, indicating NRCY is also an NLR modulator like NRCX. Furthermore, NRCX suppression of Sw5b-Nsm and NRC3-mediated cell death is dependent on NRCY. Remarkably, we found that *NRCX* and *NRCY* expressions were induced during plant senescence, while *NRCY* was induced more than *NRCX*. Accordingly, the plant resistance was stronger during maturation, indicated NRCX/Y might be involved in age-dependent resistance. Our study reveals one of the paired NLRs coordinately regulates ETI to facilitate age-dependent immunity.

## Introduction

Plants have evolved a sophisticated innate immune system to prevent infection from diverse pathogens. PRRs in the cell surface trigger plant immunity by recognizing conserved components of pathogens and hosts, known as the pattern-triggered Immunity (PTI). Successfully adapted pathogens deliver effectors to suppress defense responses including PTI. As a countermeasure, plants have evolved a second layer of immunity involving NLRs (Nucleotide-binding and Leucine-rich repeat Receptors), which recognizes pathogen effectors and trigger effector-triggered Immunity (ETI) [1,2]. ETI provides robust defense responses that are usually associated with cell death, which is referred to be hypersensitive response (HR), at infection sites to inhibit pathogen infection [3,4].

NLRs can function in pairs [5,6]. Paired NLRs are usually genetically adjacent to each other in a ‘head-to-head’ fashion and physically interact with each other [7,8]. For example, RGA4/RGA5 of rice locates in one locus in a ‘head-to-head’ form, and they physically interact with each other through CC domain regardless of the presence of effectors [9]. Usually, one of an NLR pair is a sensor, which is specialized to recognize the pathogen and the other is involved in initiating immune signaling (executor) [10]. Most executors are auto-activated and overexpression of these executors causes autoactive cell death *in planta* [10]. The autoactivity of executors can be suppressed by their sensors and knocking-out sensors may lead to lesion mimic phenotypes in plants [11,12]. The downstream signaling pathway of NLR pairs partially involves helper NLRs (NLR-Hs), e.g. ADR1 and NRG1 [13], which may form Ca^2+^-permeable cation channels to directly regulate cytoplasmic Ca^2+^ levels and consequent cell death [14].

NLRs work not only in pairs, but also through a complex network architecture [15]. Recently, a major clade of NLRs in Solanaceae plant species was shown to form a comprehensive network. In this network, multiple helper NLRs, known as NLR-Hs (Helpers NLR REQUIRED FOR CELL DEATHs), are required by a large number of sensor NLRs that mediate resistance against diverse pathogens [16]. For example, Sw5b, an NLR from *Solanum lycopersicum*, could recognize Nsm from Tospoviruses and trigger a NRC2/3-dependent HR [16,17]. Unlike typical paired NLRs, NRC-Hs are not always clustered with sensor NLRs in plant genomes even though they are evolutionarily related [16]. The current evolutionary model for the NRC network is that it has evolved from a few genetically linked NLRs, as in the Caryophyllales, through a massive expansion of sensors and a relatively limited expansion of helpers, possibly to maintain the robustness of the network against rapidly evolving pathogens [6,18].

NLRs share a basic protein architecture consisting of a variable N-terminal domain, a central nucleotide-binding domain (NB-ARC), and a C-terminal leucine-rich repeat domain (LRR) [19]. NB-ARC domains can be subdivided into three subdomains: NB, ARC1 and ARC2 [19,20]. The NB subdomain contains two major motifs, a P-loop motif required for nucleotide binding and Walker B required for adenosine triphosphate (ATP) hydrolysis [21]. In addition, there is also a nucleotide binding site MHD (methionine-histidine-aspartate) motif on ARC2 [22]. Mutation in the NLR Walker B motif or MHD motif often leads it to be auto-activated [23]. For example, the substitution of the conserved second aspartate to glutamate will reduce ATP hydrolysis rates of I-2 and lead it to autoactivation. Notably, the MHD motifs of many executor NLRs are non-canonical, and are required for auto-activation [23]. For example, the MHD motif of RGA4 is “TYG”, which is highly degenerated, differs from the consensus MHD sequence. Changing the degenerated motif to canonical MHD motif impairs the RGA4 autoactivity [9].

Age-dependent immunity is referred to the phenomenon that plant resistance to pathogens gradually increases with plant age [24]. Age-dependent immunity extensively exists in different plants. For example, *N. benthamiana* has age-dependent resistance against *P. infestans*, by which 35 days old plants have performed a completely resistance to *P. infestans* [25]. Recently, many molecular mechanisms behind age-dependent immunity have been discovered. In *N. tobaccum*, NLRs and sRNAs are reported to be associated with age-dependent immunity. During the maturation process of plants, NLRs expressions are increasing while small RNAs (sRNAs), which served as the regulatory of NLRs, are gradually decreasing in the expression [26]. The works provide a glimpse into mechanisms behind age-dependent immunity. However, other genes involved in age-dependent immunity are largely unknown.

It is proposed that functional specialization is a critical strategy in NLR evolution to avoid being constrained by signaling activity during evolution [6]. NLR networks built by helper NLR, as well as paired NLRs, require more than one NLR to mediate immune activity. We suspected that there may be redundancy between the functions of paired NLRs and helper NLRs. In this study, by analyzing the distribution of paired NLRs in Solanaceae species, we found that the quantities of paired NLRs and NRC-Hs are negatively correlated. We further used *N. benthamiana*, a model plant of Solanaceae, to evaluate the paired NLRs functions. We found that *N. benthamiana* exhibited significant dwarfing and senescence phenotypes when one of the six paired NLRs, *NRCX*, was silenced. *N. benthamiana* knockout mutant *nrcx* confirmed that the defense responses of *nrcx* are continuously activated. Interestingly, silencing the head-to-head NLR of *NRCX, NRCY*, could restore the stunting and senescence phenotype. NRCY is not a autoactive NLR but a positive modulator of NRC3-mediated immunity. Finally, we observed a gradual increase in *NRCX*/*NRCY* expressions as the leaves became senescent, where *NRCY* is induced greatly more, implied that they play a role in age-dependent resistance of *N. benthamiana*.

## Results

### The quantities of paired NLRs and NRC-Hs are negatively correlated in Solanaceae species

To evaluate the distribution of paired NLRs and NRC-Hs in Solanaceae species, we first used NLRtracker tool to identify all NLRs in 29 representative plant species including nine Solanaceae species (Fig 1A, S1 Fig). Assembled genome sizes of these organisms ranged from 89 Mb to 2,918 Mb, with annotated protein counts ranging from 23,879 to 69,500 (S2A Fig). In total, we identified 8,795 non-redundant NLR, ranging from 32 to 852 in each species (Fig 1A). The number of NLRs identified in this study is highly similar to the previous publication [27], supporting the robustness of our pipeline (S2B Fig). Paired NLRs in all these species were identified based on searching NLRs in a “head-to-head” fashion with enclosing no more than 2 non-NLR genes (Fig 1A, S1 Fig) [28]. Helper NLRs (NRC-Hs) were identified through phylogenetic assay (Fig S1, S2 File): the NLRs in the same clades with NRC2/3/4 were considered as NRC-Hs [29] (Fig 1A).

**Fig 1.**
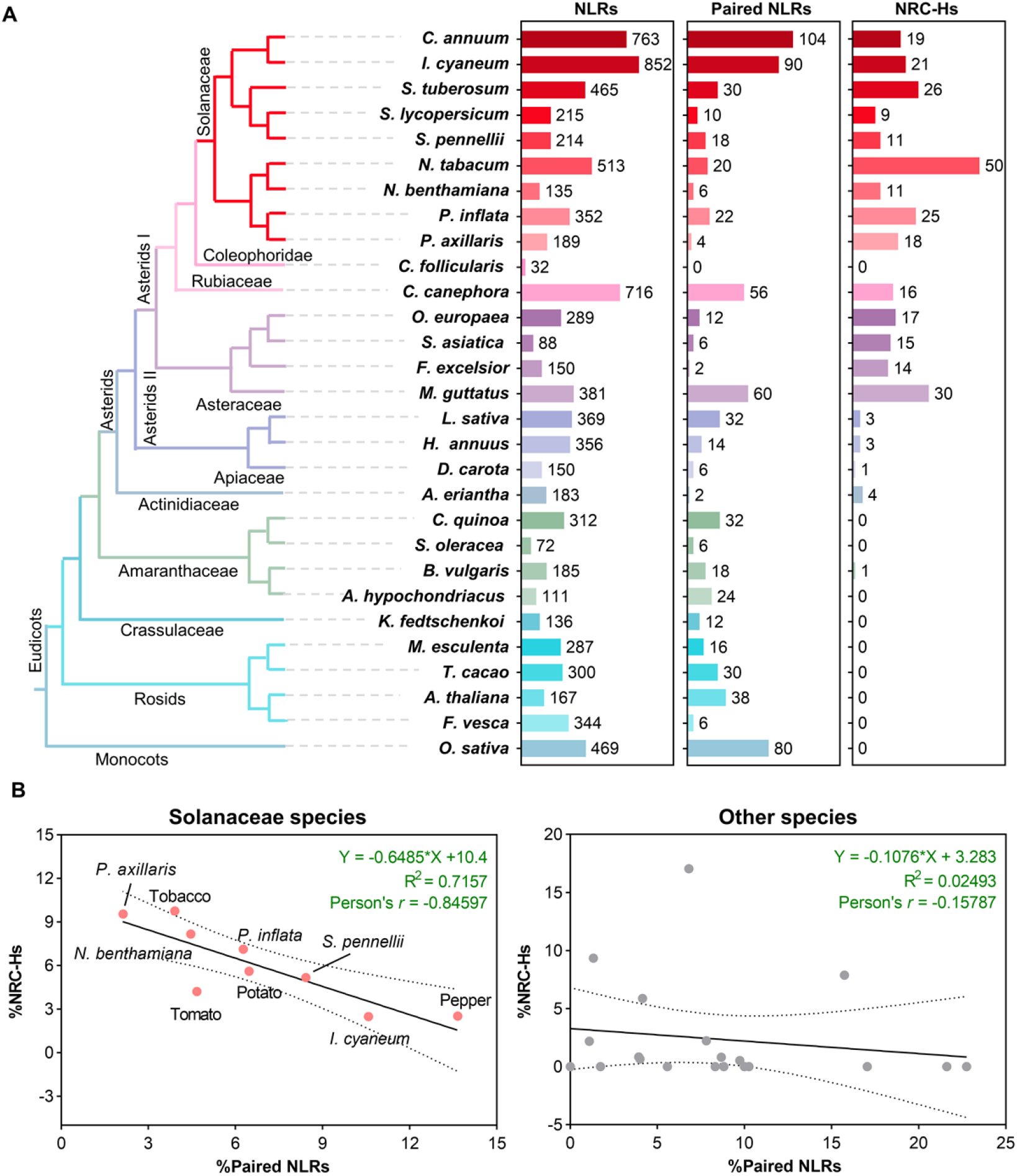
The quantities of paired NLRs and NRC-Hs are negatively correlated in Solanaceae species. (A) Summary of predicted NLRs. The phylogeny of the 29 species is based on data from the Taxonomy Database (https://www.ncbi.nlm.nih.gov/taxonomy). The counts of NLRs, paired NLRs and NRCH-s in each species are shown in boxplots following the species names. (B) Scatter plot of %Paired NLRs against %NRC-Hs. Black line represents the linear trend, with dotted line representing the 95% confidence interval. Regression equation (with its R square value) and Person correlation value is shown at top right.

We next investigated the correlation between the %Paired NLRand %NRC-Hs through regression analysis. Interestingly, the %Paired NLRs are negatively correlated with the %NRC-Hs in Solanaceae species (Fig 1B). The slope of the linear fitting equation is -0.6485 with a high R square (R^2^ = 0.7157). Furthermore, Pearson correlation analysis indicated a high negative correlation between %Paired NLR and %NRC-Hs (Pearson’s *r* = -0.84597) in Solanaceae species. In contrast, no correlation between %Paired NLRs and %NRC-Hs could be detected in other tested species (R^2^ = 0.02493, Pearson’s *r* = -0.15787) (Fig 1B). We also identified NLRs in “head-tail” or “tail-tail” fashion in Solanaceae species and found no correlation between them with NRC-Hs could be detected (S3A and S3B Figs).

### Silencing *NRCX* leads to a dwarf and accelerated senescence phenotype in *N. benthamiana*

The negative correlation between paired NLRs and NRC-Hs probably resulted from the functional redundancy between them during evolution, which might reduce the selection pressure on paired NLRs and result in loss of function or even novel function. To test it, we used the *N. benthamiana* as a model. The *N. benthamiana* only has 6 paired NLRs, which are paired NLR 1a/b (PN1a/b), PN2a/b and PN3a/b (Fig 2A). To evaluate their functions, we silenced all 6 paired NLRs individually (Fig 2A). We observed a different phenotype only when *PN3b* was silenced (Fig 2B). qRT-PCR analysis indicated that all the paired NLRs were silenced successfully (S4A Fig).

**Fig 2.**
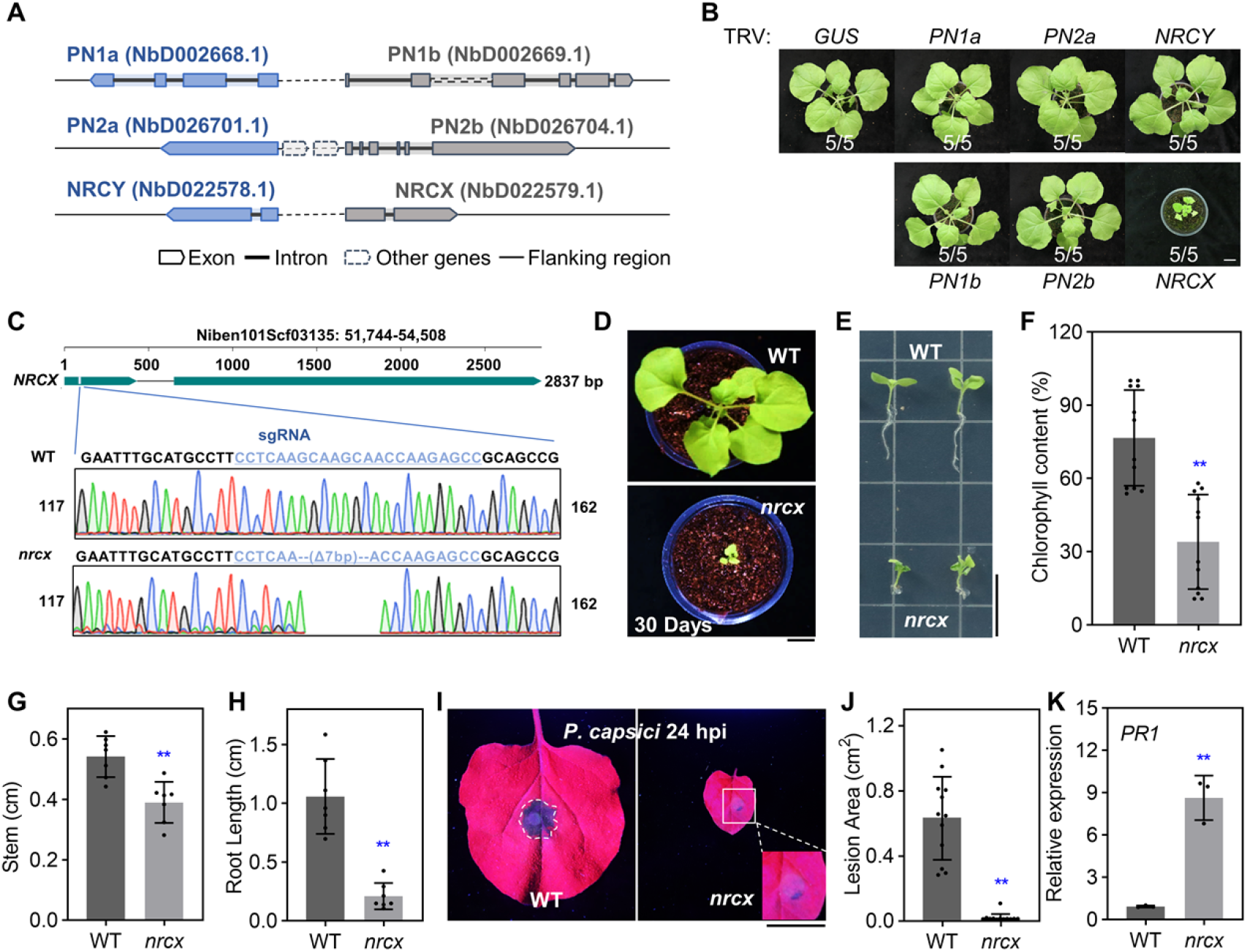
Mutant of *NRCX* is senescence-accelerated, display growth defect and high resistance level. (A) Schematic diagram of six paired NLRs in *N. benthamiana*. (B) The morphology of plants silencing indicated genes. Two-week-old *N. benthamiana* plants were infiltrated with *Agrobacterium* strains carrying tobacco rattle virus (TRV) VIGS constructs, and photographs were taken 4 weeks after agroinfiltration. TRV:*GUS* (*β-glucuronidase*) was used as a negative control. Bar = 2 cm. (C) Schematic diagram showing the deletion of *NRCX* in *N. benthamiana*. The genomic region information of *NRCX* is shown above. The gRNA used for CRISPR/Cas9 editing and DNA sequencing results are shown below. (D) Phenotypes of wild type and *nrcx* knockout mutants grown in soil for 30 days. Bar = 2 cm. (E) Phenotypes of wild type and *nrcx* mutant seedlings on sterile 1/2 MS medium for three weeks. Bar = 1 cm. (F) Chlorophyll content in leaves of *N. benthamiana* wild type and *nrcx* mutant. Asterisks indicate statistically significant differences with t test (mean ± SD, n = 12, \*\**P* < 0.01), and asterisk in the following text indicates the same meaning. (G,H) Stem length and root length statistics of wild type and *nrcx* grown on 1/2 MS medium for 3 weeks (mean ± SD, n = 7). (I-J) Enhanced resistance in *nrx* mutant. The photos were taken at 24 hours after inoculation (hpi) under UV light. Bar = 2 Inoculated lesion areas are show in (J) (mean ± SD, n = 12). (K) Upregulation of *PR1* expression in *nrcx* mutant. The indicated leaves were inoculated with *P. capsici* for 12 hours and the *PR1* expression was tested (mean ± SEM, n = 3).

We then focus on the function of PN3b. Further sequence analysis indicated that PN3b is NRCX, an NLR modulator protein negatively regulates NRC2/3-mediated immunity response identified in a parallel research [30]. In addition to a dwarf phenotype (S4B Fig), silencing *NRCX* also made the plant senescence accelerated (S4C and S4D Figs). The tobacco rattle virus accumulation levels in NRCX-silenced plants were lower than those in control lines, indicating that the dwarf phenotype is not resulted from over-accumulated virus (S4E Fig).

### The senescence-accelerated and dwarf of *nrcx* mutant are spontaneous

To further remove the possibility that such phenotype was caused by the virus or off-target effect of VIGS technology, we used CRISPR/Cas9 system to generate *NRCX* loss-of-function mutants. One small guide RNA (sgRNA) was designed based on the unique sequence of NRCX (Fig 2C) and transformed together with Cas9 and a kanamycin selection maker into *N. benthamiana*. Nine independent transformants were selected using kanamycin. Among the recovered lines, only one transformant containing a 7 bp deletion in the *NRCX* locus was a homozygous *nrcx* mutant (Fig 2C), while the other transformants were heterozygous or wild-type. No off-target effects were found in *NRCX* homologues (S5A and S5B Figs). The development of *nrcx* mutants was significantly arrested compared to that of the wild type (Fig 2D), suggesting that loss-of-function of *NRCX*, rather than the virus and off-target effect of VIGS technology, was responsible for the phenotypes.

To figure out whether the *nrcx* phenotypes are spontaneous or stimulated by the microbiome in the soil, *nrcx* and wild-type plants were grown in microbiome-free medium (Fig 2E). Compared with wild type, *nrcx* mutants had a high degree of chlorosis and were smaller (Figs 2F-H), indicating that the dwarfing and accelerated aging phenotype of *nrcx* is spontaneous. Importantly, we found that *nrcx* mutants showed higher inhibition levels in the root than in the stem (Figs 2G and 2H), indicating that NRCX plays a vital role in root development, which is consistent with the report that *NRCX* is highly accumulated in roots [30].

### Defense response is activated in *nrcx* mutant

Plant growth and resistance are often in a state of ebb and flow, the dwarf and accelerated senescence phenotype may result from the persistent activation of plant immunity [31-33]. *Phytophthora* inoculation assay indicated that *nrcx* plants exhibited high levels of resistance to *P. capsici* (Figs 2I and 2J). To test whether the defense response was activated in *nrcx* mutants, the *PR1* (*Pathogenesis-related protein 1*) gene of wild-type and *nrcx* mutants were tested separately, and *PR1* was up-regulated about ∼10 fold in *nrcx* mutant plants (Fig 2K). Collectively, these results suggest that defense responses are continuously activated in *nrcx* plants.

### Simultaneously silencing *NRCY* restores the *NRCX*-silenced plants phenotype

Knocking-out one paired NLR may activate their conjugated NLR to activate defense pathways and inhibit plant growth [34]. The head-to-head NLR (*PN3a*) of *NRCX* is named after *NRCY* (*NLR-REQUIRED FOR CELL DEATH Y*) (Fig 2A). *NRCY* and *NRCX* are tandemly linked in a head-to-head manner on a scaffold with an intergenic region of 18,795 bp between them. We assumed that, like other NLR pairs, *NRCX* silencing may activate NRCY and subsequently induce plant defense responses. To verify this, we co-silencing *NRCX* and *NRCY* in plants using one TRV vector where the gene fragments of *NRCX* and *NRCY* were tandemly cloned. Compared to TRV:*NRCX* plants, growth inhibition and accelerated senescence are significantly restored in TRV:*NRCX/Y* plants, even though TRV:*NRCX/Y* plants were still smaller than TRV:GUS (*β-glucuronidase*) plants (Figs 3A-C). qRT-PCR assays indicated that these genes had been successfully silenced (Figs S6A and S6B).

**Fig 3.**
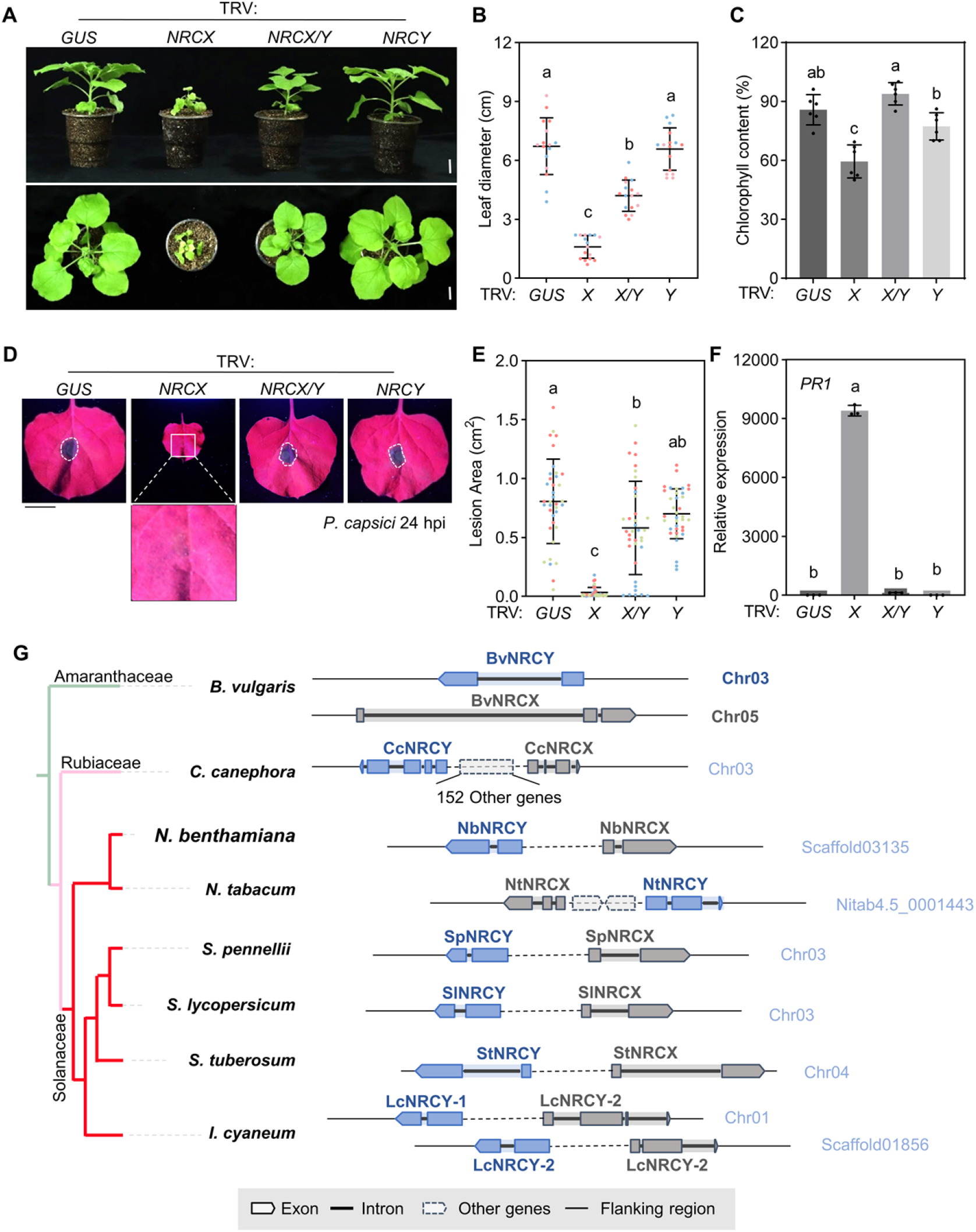
Simultaneously silencing *NRCY* restores the *NRCX*-silenced plants phenotype. (A) The morphology of *N. benthamiana* plants silencing indicated genes. TRV:*GUS* was used as a negative control. Two-week-old *N. benthamiana* plants were infiltrated with *Agrobacterium* strains carrying TRV constructs, and the photographs were taken 4 weeks after agroinfiltration. Bar = 2 cm. (B) Leaf diameter of plants silencing indicated genes. The fourth and fifth true leaves (counting from bottom to top) were harvested and the average diameters were recorded. Three independent biological replicates were performed and indicated by different colored dots (n = 16, lowercase letters indicate significant differences tested between multiple groups by one-way ANOVA at P < 0.05). (C) Chlorophyll contents of indicated leaves. The fourth true leaf (counting from bottom to top) was taken from five plants silencing indicated genes at 6 weeks of age and the chlorophyll contents were calculated (n = 6, lowercase letters indicate significant differences tested between multiple groups by one-way ANOVA at *P* < 0.05). (D-E) *P. capsici* inoculation phenotypes. Leaves of plants silencing indicated genes were detached and inoculated with *P. capsici* and then incubated in a growth room at 25°C in darkness. Bar = 2 cm. Lesion area was calculated and is shown at (E). Different colored dots indicate different repetitions (n = 37, lowercase letters indicate significant differences tested between multiple groups by one-way ANOVA at *P* < 0.05). (F) Relative *PR1* gene expression levels. The indicated leaves were inoculated with *P. capsici* for 12 hours and the PR1 expression was tested (mean ± SEM, n = 3). (G) Schematic diagram of NRCX and NRCY plant genomes.

We next tested whether the activated immunity in TRV:*NRCX* plants is impaired in TRV:*NRCX/Y*. The enhanced resistance to *P. capsici* in TRV:*NRCX* plants was partially restored in TRV:*NRCX/Y* (Figs 3D and 3E). Similarly, the *PR1* expressions in *NRCX/Y*-silenced plants were significantly lower than that in TRV:*NRCX* plants (Fig 3F). Collectively, these results indicated that NRCY was a positive regulator of plant immunity and senescence. And the dwarf and accelerated senescence phenotype of *NRCX*-silenced plants is partially dependent on NRCY.

### NRCX/Y pair is specific in Solanaceae species

We further tested whether the NRCX and NRCY conserved. According to our phylogenetic assay (S7 Fig). NRCX homologs could be first discovered in *B. vulgaris* during evolution and are specific in Asterids species, which is consistent with the previous results [30]. Meanwhile, NRCY are conserved in dicots (S7 Fig). We further analyzed the distribution of paired NRCX/Y by analyzing whether they are tandemly located in one locus (no more than 2 other genes between them). In *B. vulgaris* (sugar beet), NRCX and NRCY homologs are located in different chromosomes (Fig 3G). In *C. canephora* (coffee), NRCX and NRCY are found in same chromosomes but separated by 152 other genes (Fig 3G). In contrast, NRCX/Y are adjacent to each other and conserved in *N. tabacum, N. benthamiana, S. pennellii, S. lycopersicum, S. tuberosum, I. cyaneum*, indicated that the paired NRCX/Y is specifically encoded by Solanaceae species (Fig 3G).

### The walker B and MHD motifs of NRCY are non-canonical

*NRCY* encoded an 881-aa CNL containing a CC domain, an NB-ARC domain and an LRR domain. P-loop, walker B and MHD motifs within the NB-ARC domain are important for NLR activation and mutations of critical residues in these motifs may lead to NLR autoactivation [23]. Sequence analysis indicated that the P-loop sequence of NRCY matches the consensus sequence (GxxxxGK[T/S]) (Fig 4A). Interestingly, in the walker B motif of NRCY, the conserved second aspartate is naturally substituted by glutamate (Fig 4A), and this mutation was reported to result in reduced ATP hydrolysis rates and autoactivation of I-2 [35]. Furthermore, the MHD motif of NRCY is highly degenerated, with all three amino acids distinct from the consensus MHD sequence (Fig 4A). Similarly, NLRs with the non-canonical MHD motif also frequently induce cell death [23].

**Fig 4.**
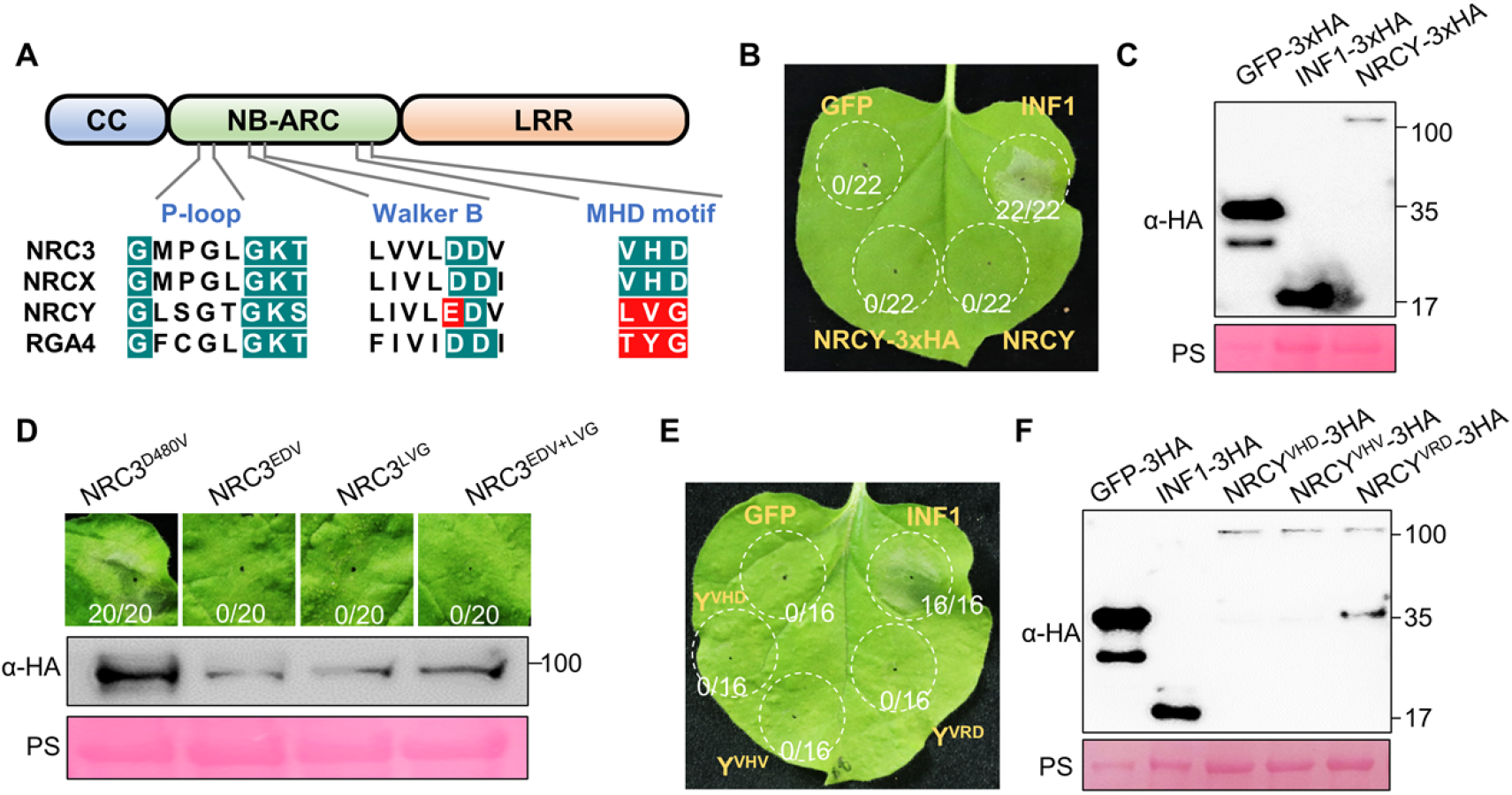
NRCY does not induce HR in *N. benthamiana*. (A) P-loop, walker B and MHD motifs of indicated NLRs. The motif sequences of NRC3, NRCX, NRCY and RGA4 are shown. Canonical residues in motifs are marked by blue while non-canonical residues are marked by red. (B) The phenotype of leaves expressing NRCY. GFP and INF1 were used as negative and positive controls for cell death, respectively. (C) Protein accumulation of NRCY in *N. benthamiana*. Proteins were extracted from leaves at 36 hours after the agroinfiltration. Indicated protein was detected using HA antibody. Total protein loading was confirmed by Ponceau S staining. (D) Cell death induced by the NRC3 mutants. NRC3^D480V^ is an autoactive NRC3 mutant. The phenotypes of leaves expressing NRC3 with walker B motif of NRCY (NRC3^EDV^), NRC3 with MHD motif of NRCY (NRC3^LVG^) and NRC3 with both walker B and MHD motif of NRCY (NRC3^EDV+LVG^) are shown above. Accumulation of the NRC3 variant proteins is shown below. (E) Phenotype of leaves expressing autoactive NRCY mutants. Photos were taken at 5 days post infiltration. (F) Protein accumulation of NRCY autoactive mutants. Total protein loading was confirmed by Ponceau S staining.

Phylogenetic assay showed that the walker B motif and MHD motif of all NRCY homologs in Arabidopsis, kiwifruit and sugar beet and are canonical (S7 Fig). The non-canonical MHD motif emerged in partially NRCY homologs of coffee, while non-canonical Walker B motif emerged in Solanaceae species (S7 Fig), indicating that these two motifs might degenerate during the NLR evolution in Solanaceae species.

### NRCY does not induce autoactive cell death in *N. benthamiana*

Based on the non-canonical walker B and MHD motifs in NRCY, it is assumed that NRCY is an automatically activated NLR. Transient expression was performed on *N. benthamiana* to detect whether NRCY was self-activating according to the phenomenon of cell death. Surprisingly, no cell death was detected when expressing *NRCY* in leaves, regardless of whether the protein was fused to a tag (Fig 4B). Western blot analysis indicated that *NRCY* was expressed correctly (Fig 4C).

These results inspired us to investigate whether the walker B and MHD motifs of NRCY are auto-activated mutants. The original walker B and MHD motifs of NRC3 were replaced by the walker B and MHD motifs of NRCY. NRC3 with walker B motif of NRCY (NRC3^EDV^), NRC3 with MHD motif of NRCY (NRC3^LVG^) and NRC3 with both walker B and MHD motifs of NRCY (NRC3^EDV+LVG^) induced no cell death when overexpressed in *N. benthamiana*, while NRC3^D480V^ triggered severe cell death (Fig 4D). The protein levels of NRC3^EDV^, NRC3^LVG^ and NRC3^EDV+LVG^ were lower than NRC3^D480V^, but these mutants were expressed correctly (Fig 4D). These results indicated that the walker B and MHD motif of NRCY were not auto-activated mutants. Next, we changed the mutated MHD motif of NRCY to typical auto-activating mutants (NRCY^VHV^ and NRCY^VRD^) and they also triggered no cell death (Figs 4E and 4F). These results suggest that, like NRCX [30], NRCY does not possess the ability to induce cell death.

### Silencing *NRCY* impairs Nsm-Sw5b and NRC3-mediated cell death

The observation that NRCY is a genetic suppressor of NRCX and is not an auto-activated NLR leads us to ask whether NRCY is another NLR modulator that regulates the function of NRC network. To test this, RNAi technology was used to silence *NRCY* in *N. benthamiana* leaves (S6C Fig), and Sw5b/Nsm, Avr3a/R3a, auto-activating mutants of NRC3 and NRC4 were subsequently expressed. No differences in cell death mediated by NRC4 auto-activating mutants, Avr3a/R3a and INF1 was detected in *NRCY*-silencing leaves (Fig 5A). However, *NRCY*-silencing led to impaired cell death induced by NRC3^D480V^ and Nsm-Sw5b (Figs 5A and 5B), which indicated that NRCY is a positive modulator of NRC3 function.

**Fig 5.**
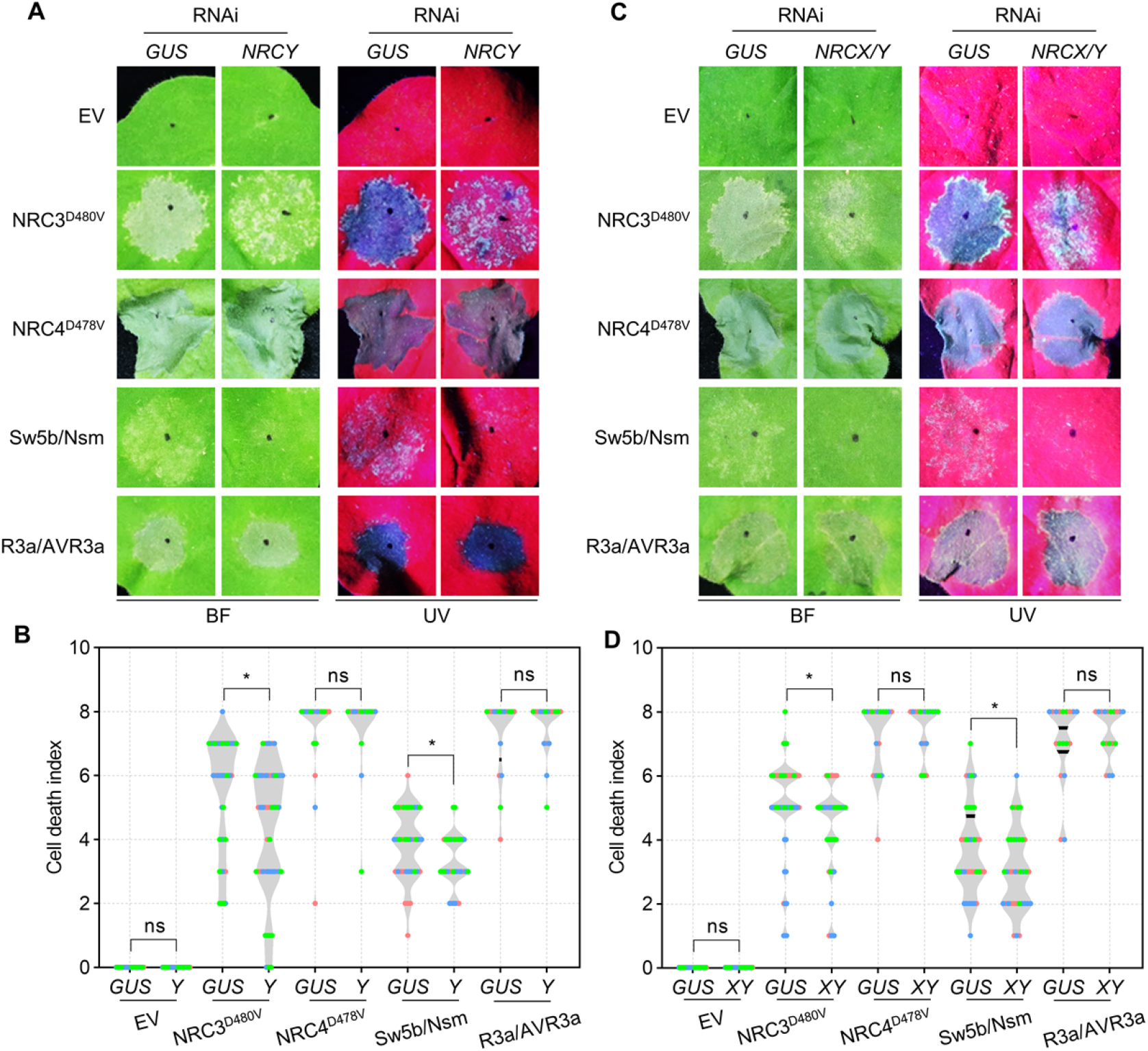
Silencing NRCY impairs Nsm-Sw5b and NRC3-mediated cell death. (A-D) Hypersensitive response phenotypes in indicated leaves. Photos were taken at 5 days post infiltration and are shown (A,C). Violin plots in (B,D) showing the HR index scored at 5 days post agroinfiltration. Three independent biological replicates were performed and indicated by different colored dot. Asterisks indicate statistically significant differences with *t* test (n > 14, \*\**P* < 0.01).

To characterize the genetic relationship between NRCX and NRCY, *NRCX* and *NRCY* were co-silenced. Although silencing *NRCX* enhanced cell death induced by NRC3^D480V^ and Nsm-Sw5b (S8A-C Figs) [30], co-silencing *NRCX* and *NRCY* still inhibited the cell death, consistent with phenotypes in plants silencing *NRCY* alone (Figs 5C and 5D). These results indicated that the function of NRCX in regulating NRC3 is dependent on NRCY. Collectively, NRCX/Y are paired NLR modulators that coordinately regulate NRC3-mediated immunity.

### Association occurs between CC domains of NRCY and NRCX

Paired NLRs are adjacent to each other at the genome locus and often display physical interactions [6,36]. As our data indicated that NRCX/Y is not a typical NLR pair, it is interesting to test whether there is still a physical interplay between NRCX and NRCY. However, we could not express full-length NRCX protein in *N. benthamiana* correctly, so we did not investigate their association using full length NRCX/Y. Instead of that, we test the association between their domains via split-luciferase assays and yeast two-hybrid (Y2H) (S9A Fig). The results indicated that the CC domains of NRCY and NRCX interact with each other (S9B and S9C Figs).

### *NRCX* and *NRCY* are involved in age-dependent plant immunity

Head-to-head NLR pairs are often transcribed at an equal rate to prevent autoimmune triggered by the executors. As NRCX/Y are NLR modulators, we asked whether the transcriptions of *NRCX* and *NRCY* are still synchronous. Interestingly, when we analyzed the expression of *NRCX* and *NRCY* during leaf senescence, we found that, although the expression of *NRCX* and *NRCY* increased gradually during leaf senescence, their growth rates were different (Figs 7A and 7B). *NRCX* was up-regulated 5-fold in mature leaves compared to young leaves, whereas *NRCY* induced ∼30-fold change (Figs 7A and 7B).

**Figure 7.**
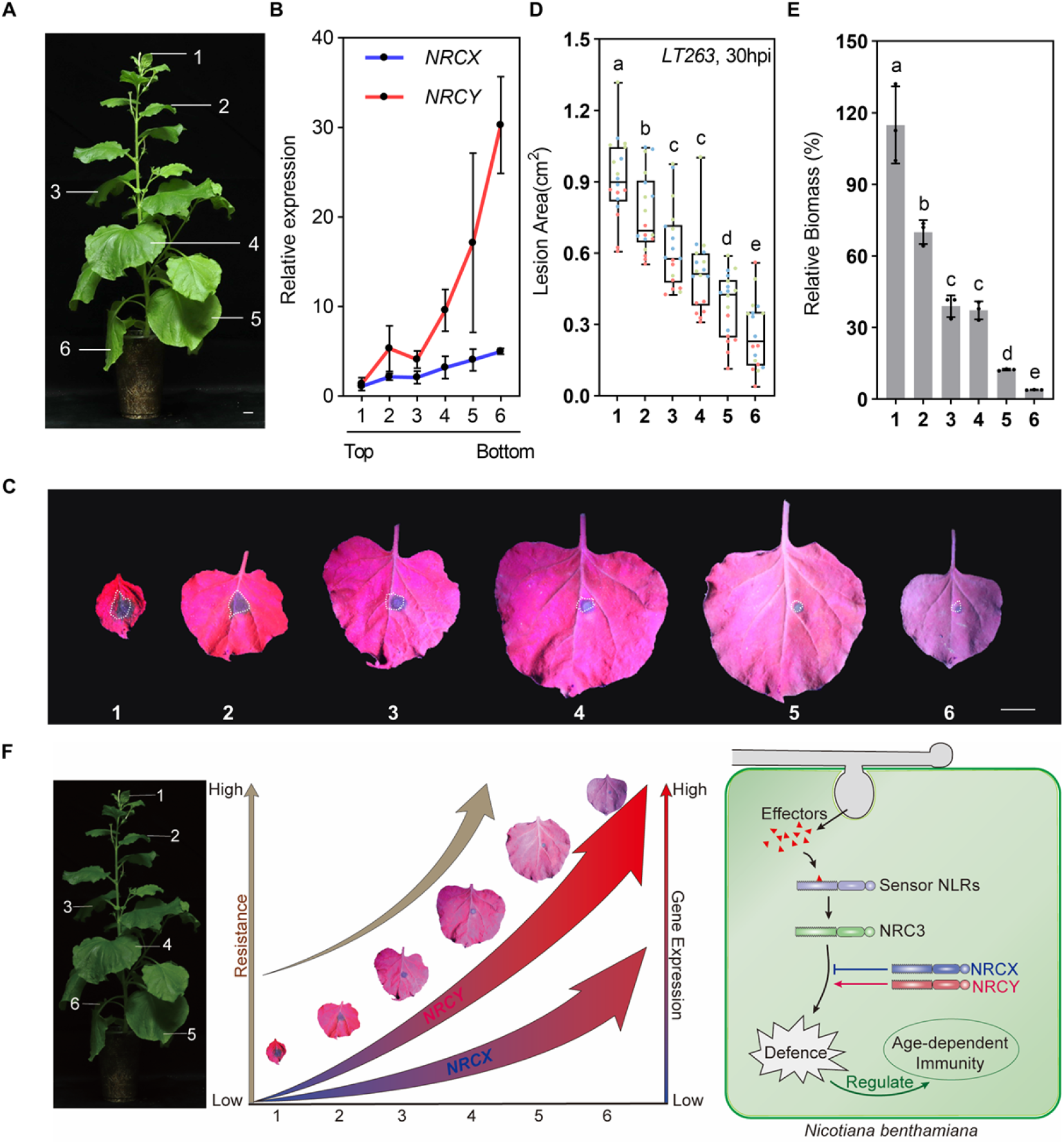
*NRCX* and *NRCY* are involved in age-dependent plant immunity. **(A**,**B) Upregulation of *NRCX* and *NRCY* during leaf senescence**. Leaf positions used to evaluate *NRCX/Y* expressions are labelled in (A). A total of 6 leaves in each positions were sampled for qRT-PCR assay. The relative expression levels of *NRCX* and *NRCY* in leave 1-6 are shown in (B) (mean ± SEM, n = 3). Bar = 2 cm **(C-E) Age-dependent immunity of *N. benthamiana to P. capsici***. Leaves from positions indicated in (A) were sample and inoculated with *P. capsici* mycelium. The photos were taken at 24 hpi (C). Lesion areas were calculated and are shown in (D). Three independent replicates were performed and labelled by different colors (mean ± SD, n > 15, lowercase letters indicate significant differences tested between multiple groups by one-way ANOVA at *P* < 0.05). Relative biomass of *P. capsici* is shown in (E) (mean ± SEM, n = 3, lowercase letters indicate significant differences tested between multiple groups by one-way ANOVA at *P* < 0.05). (F) illustration of working model. The paired NLRs, NRCX/Y, coordinately modulate NRC3-dependent immunity and are involved in age-dependent immunity through different expression level during leave maturation.

Age-dependent immunity is referred to the phenomenon that the plant resistance to pathogens is gradually increases during plant maturation [24]. We tested the resistance of senescent leaves and young leaves to *P. capsici*, and found the resistance increased gradually (Figs 7C-E). The high growth rate of *NRCY* expression level and the relatively low growth rate of *NRCX* expression level during leaf senescence indicated that the paired NLR modulators may be involved in age-dependent immunity (Fig 7F).

## Discussion

We identified the paired NLRs and NRC-Hs in the representative Solanaceae species and found a negative correlation between their numbers, suggestive of their potential redundant functions. Using a knock-down assay, we found that *NRCX*-silenced *N. benthamiana* lines exhibited obvious dwarfism and accelerated senescence, which could be restored when its head-to-head NLR pair, *NRCY*, was silenced. Furthermore, we demonstrated that *NRCX* and *NRCY* differently modulates NRC3-mediated immunity, thereby coordinately regulating ETI response in *N. benthamiana*. Finally, we found that *NRCX* and *NRCY* were up-regulated during plant senescence, while the expression of *NRCY* was induced more than *NRCX*. Accordingly, plant resistance was getting stronger throughout maturation, indicating NRCX/Y might be involved in age-dependent resistance. Collectively, we propose that the paired NLRs, NRCX/Y, are novel modulators in age-dependent immunity by coordinately regulating NRC3-mediated immunity (Fig 7F).

Functional specialization is a vital event in NLR evolution that enhances NLRs capacity to keep up with rapidly evolving pathogens without being constrained by signaling activity [6]. In Solanaceae species, NRC-Hs and their sensors expanded greatly [16]. The putative redundant function of NRC-Hs and paired NLRs might reduce the selection pressure of paired NLRs. In this study, we identified the paired NLRs in Solanaceae species and found a high negative correlation between paired NLRs and NRC-Hs. Further silencing experiments showed that, unlike a parallel work in rice, when one of the putative NLR pairs was knocked out, the lesion-mimicking or dwarfing phenotype was widespread [28]. Moreover, we found that *N. benthamiana* exhibited significant dwarfing and senescence phenotypes only if *NRCX* was silenced, which is consistent with the previously study [30]. Furthermore, although NRCX/Y contain many feathers of typical paired NLR, they are NLR modulators. These results might indicate that paired NLRs have degenerated during NRC-Hs expansion in Solanaceae species. Interestingly, in pepper and Iochroma, the situation is reversed: %NRC-Hs is relative fewer and the %paired NLRs are large in number, indicated that NLR pairs still function majorly in ETI in the two species.

Paired NLRs work together to regulate immunity and usually play different functions. One of them specifically recognizes effectors through specific domains such as HAM (Heavy-Metal Associated domain) and WRKY, and the other activates immune response including cell death [37-40]. However, the NLR pair NRCX/Y found in this study acts more like immune modulators. Neither NRCX nor NRCY has obtained an integrated domain that can recognize effector structures like HAM and WRKY domain. Moreover, multiple putative activating mutants of NRCX and NRCY could not induce the HR response of *N. benthamiana* [30], indicating that the mode of action of NRCX and NRCY is different from the traditionally considered paired NLRs. Instead of that, the NRCX/Y modulate NRC3-dependent immunity. On the other hand, NRCX/Y still maintain some typical feathers of NLR pair. Firstly, they located in genome in a “head-to-head” fashion. Secondly, knocking out *NRCX* did induce a dwarf phenotype, which is reminiscent of knocking out sensor NLR in a typical NLR pair. Thirdly, paired NLRs usually interact with each other [9,40,41], and we found their CC domains can form heterodimer. These indicated that the NRCX/Y pair might be in a transitional stage with both original and novel feathers.

The NB-ARC domain of NLRs acts as a molecular switch for ATP hydrolysis, switching from the ‘off’ state of ADP binding to the ‘on’ state of ATP-binding to activate downstream immune responses [21,42]. The walker B motif (hhhDD/E) and MHD motif motifs are important for ATP hydrolysis and ADP binding, respectively [22,43,44]. Mutation in the two motifs often results in an autoactive phenotypes [45]. Although NRCY has a non-canonical walker B (LIVLEDV) and MHD motif (LVG), it cannot induce cell death. The results further supported that NRCY is more of a modulator than an executor. Interestingly, we found the non-canonical motifs of NRCY emerged in coffee, which is closely to the emergence of paired NRCX/Y specific in Solanaceae species. It is speculated that NRCY might be degenerated from a functional NLR and display a genetically linkage with NRCX after the split between Solanaceae species and Coleophoridae species.

*NRCY*-silencing lead to impaired cell death induced by NRC3^D480V^ and Nsm-Sw5b. Furthermore, co-silencing of NRCX and NRCY and NRCY alone consistently both inhibited cell death, suggesting that NRCX regulation of NRC3 function is dependent on NRCY. However, it remains unknown how paired NLRs regulate NRC3-mediated cell death. Many NLRs induce cell death by forming resistosomes. For example, CNLs are activated to form pentamers with cation channels, while TNLs are activated to form tetramers [46-49]. RNLs belong to helper NLR downstream of TNL immunity, which form pentamers in calcium ion channels and finally induce immune responses such as cell death in plants [14]. Recently, NRC2 from NRC-H clade oligomerizes upon effector detection by its sensor NLR [50], indicating that NRC-Hs also form resistosomes. A hypothesis is that NRCX/Y pair might regulate the resistosome-forming process and thus effect the final cell death. It is interesting to determine the mechanism how NRCX/Y regulating the function of NRC-H proteins in the future.

Our data expands the current definition of NLR modulators. Two NLR modulators, NRCX and NRG1C are reported and both of them are helper NLRs [30,51]. NRCX belongs to NRC-H family. It contained all NRC-Hs feathers including the MADA motif in N terminal. However, NRCX does not induce cell death but acts as a modulator [30]. Overexpression of NRG1C suppresses autoimmunity of NRG1A, *snc1* and *chs3-2D*, without affecting *chs1, chs2* autoimmunity and RPS2-and RPS4-mediated immunity [14]. Intriguingly, unlike NRCX, NRG1C is a truncated NLR that lacking the whole N-terminal CC_R_ domain and is therefore unlikely to execute the hypersensitive cell death [51]. Here we show that NRCY, which is the conjugated NLR pair of NRCX, is also an NLR modulator. Furthermore, the suppression of NRCX in NRC3 is dependent on NRCY, indicating that NRCX and NRCY are paired NLR modulators.

In eukaryotic genomes, there are approximately 10% of genes are arranged in a head-to-head orientation. These head-to-head orientation often tend to be genes that should be transcribed at equal rates for gene products, suggesting a pattern of shared regulatory regions [52]. For example, co-expression from a head-to-head cluster has been observed for the SOC3– CHS1–TN2 NLR cluster in Arabidopsis, in which the gene products do interact [53,54]. A genome-wide analysis using microarray data for other Arabidopsis head-to-head paired NLRs further supports the co-expression pattern of paired NLRs [53]. In this study, we also observed a common tendency shared by NRCX/Y expressions. In humans, there have even been bidirectional promoters found for paired genes [55]. Although so far no NLRs have been shown to possess bidirectional promoters in plant, it is a potential reason for the co-expression of *NRCX* and *NRCY*. Interestingly, *NRCY* is up-regulated greatly more during leave senescence. We hypothesize that it is probably due to that NRCY is not auto-active and thus releases the selection pressure on restricting *NRCY* transcription during evolution. The mechanism behind should be studied in the future. Additionally, the difference in expression pattern could be used as a clue to identify novel paired NLR modulators.

High immunity level often antagonizes normal plant growth so that age-dependent immunity is a high-efficient strategy for the plant to balance this fitness cost. *N. benthamiana* is reported to have age-dependent resistance against *P. infestans* [25]. Here we showed the difference in resistance between senescent and young leaves on a plant. The senescent leaves are nearly resistant to *P. capsici*, while the young leaves are susceptive (Fig 7C-E). The age-dependent immunity is reported to be related to NLRs in the transcripts level. In *Nicotiana tabacum*, NLRs expression gradually increases with plants matured to form age-dependent immunity [26]. In this study, the expression of *NRCX* and *NRCY* also increased gradually during leaf senescence. Furthermore, the growth rate of *NRCY* expression was higher than that of *NRCX* (Fig 7A). These data combined with the findings that NRCX/Y are paired NLR modulators, together suggested NRCX/Y are involved in age-dependent immunity. In young tissues, where NRCY is less accumulated, it will facilitate plant development, while in mature tissues, highly induced *NRCY* improves plant resistance to pathogens.

Collectively, our findings show the negative correlation between paired NLRs and NRC-Hs in Solanaceae species. It highlights a previously neglected co-evolution between NRC-Hs and paired NLRs in Solanaceae species. It also shows the importance of novel role of paired NLRs as modulators in ETI and other biological processes such as in age-dependent immunity.

## Methods

### Identification of NLRs, paired NLRs and NRC-Hs

To identify NLRs, protein sequences from 29 plants were used as input for NLRtracker software [56]. The proteins containing NB-ARC (N) domain predicted by NLRtrack are NLRs. Paired NLRs in all these species were identified based on searching NLRs in a “head-to-head” fashion with enclosing no more than 2 non-NLR genes. Helper NLRs (NRC-Hs) were identified through phylogenetic assay: the NLRs in the same clades with NRC2/3/4 were considered as NRC-Hs.

### Growth Conditions of *Nicotiana benthamiana*

*N. benthamiana* plants used in this study were grown in the greenhouse at a temperature of 25°C under a 16-h light/8-h dark photoperiod and 60% relative humidity. The VIGS-treated *N. benthamiana* plants used in this study were grown in a 22°C greenhouse under a 16-h light/8-h dark photoperiod and 60% relative humidity.

### Virus-induced gene silencing (VIGS)

VIGS experiments were carried out in *N. benthamiana* as previously described [57]. The binary constructs pTRV2 were transformed into *Agrobacterium* strain GV3101-pMP90. One days before the *Agrobacterium* infiltration, the constructs were incubated at 30°C and 220 rpm for 24 hours. *Agrobacterium* were collected by centrifugation at 4000 rpm for 4 min, washed and re-suspended in infiltration buffer [10 mM MgCl_2_, 10 mM MES (pH 5.7) and 200 μM acetosyringone], the OD_600_ was adjusted to 0.3. Two-week-old *N. benthamiana* plants were infiltrated with a suspension of Agrobacterium carrying TRV RNA1 and TRV RNA2 in a 1:1 ratio.

### Transient gene expression and cell death assays

The binary expression plasmids were transferred by electroporation into *Agrobacterium* strain GV3101. Four-weeks old *N. benthamiana* were used for transient expression by *Agrobacterium*. The *Agrobacterium* suspension was configured in the same way as in the VIGS experiment above.

To perform RNAi experiments in *N. benthamiana* leaves, *Agrobacterium* carrying RNAi plasmids were injected (OD_600_=0.2) into leaves one day before transient expression of other genes. The OD_600_ of the target gene was adjusted to 0.6. The cell death phenotype was scored and photos were taken 7-days post *Agrobacterium* infiltration (dpi). HR scores were modified as from 0 (no visible necrosis) to 8 (fully confluent necrosis) (S8C Fig).

### Plasmid constructions

All plasmids and primers for the recombinant constructs used in this work are listed in S1 File. NRCX and NRCY was amplified from *N. benthamiana* cDNA and cloned into different vectors such as pCambia1300-3xHA and pCambia1300-3xFlag. Sequences of primers used in cloning of NRCX, NRCY, NRCY variants and NRC3 variants are listed in S1 File. Functional analyses of NRCY and NRC3 were performed with untagged variants, while C-terminally HA or Flag tagged variants showed consistent results with untagged variants in complementation assays.

### Measurement of Chorophyll Contents

The chlorophyll content of the fourth true leaf (counted from bottom to top) of each *N. benthamiana* plant was determined using a SPAD Chlorophyll Meter (SPAD-YA, Huoerd). Each leaf was evenly divided into 4 spots, and the average value of the 4 measurements (SPAD Unit) represents a single data point. At least four individual leaves of each treatment are measured, and three biological replicates were performed.

### RT-qPCR analysis

Plant total RNA was extracted using an RNA-simple Total RNA Kit (Tiangen Biotech Co., Ltd., Beijing, China). DNA contamination in the RNA sample was removed by 4xgDNA wiper (Vazyme Biotech Co., Ltd., Nanjing, China). 1μg of each RNA sample was subject to first strand cDNA synthesis using the HiScript II Q RT SuperMix for qPCR (Vazyme Biotech Co., Ltd., Nanjing, China). Real-time PCR was performed on an ABI Prism 7500 Fast Real-Time PCR System using the SYBR Premix Ex Taq kit (Takara Bio Inc., Shiga, Japan) according to the instructions. Gene expression levels were normalized to the expression of *NbEF1a*, which is stably expressed reference gene in *N. benthamiana*. The primers used in the RT-PCR are listed in S1 File.

### *P. capsici* culture conditions *and* inoculation assays

The *P. capsici* strain *LT263* used in this study was cultured and maintained at 25°C in the dark on 10% (v/v) V8 agar plates. The steps of inoculation experiment were as follows: mycelium blocks were placed on the back of the leaves, and 0.1% tween 20 was added at the intersection of mycelium blocks and leaves. After fully moisturizing, the mycelium blocks were placed in a 25°C incubator in darkness for 24 to 36 hours. incubated in an incubator at 25°C for 24 to 36 well moisturized and then incubated in a 25°C incubator for 24 to 36 hours, photos were taken under ultraviolet light, and the damaged area was measured by Image J software.

### Determination of biomass of *Phytophthora capsicum*

Diseased leaves of the same quality after inoculation with *Phytophthora capsicum* should be taken, and samples should include all diseased spots. The internal reference genes of *Phytophthora capsicum* and the internal reference genes of *N. benthamiana* were used as the biomass representative genes of *Phytophthora capsicum* and *N*.*benthaminan* respectively. *T*he biomass of *P. capsicum* in *N. benthamiana* was detected according to the expression levels of these two quantitative reference genes.

### Cultivation of *N. benthamiana* seedlings on 1/2 MS medium

Put 60 tobacco seeds into a 1.5 mL EP tube, sterilize with 75% ethanol for 30 s, and wash 3 times with sterile water. Then disinfect with 2.5% sodium hypochlorite for 8 minutes and rinse with sterile water 6 times. Seeds were evenly distributed on 1/2 MS medium (0.8% agar powder, pH 5.7). The medium was sealed and placed horizontally at 4 °C for 3 days to break the seed dormancy, then cultured in the greenhouse to wait for the seeds to germinate and grow, and photographed after two or three weeks.

### Protein extraction of *N. benthamiana* leaves

For normal western blot assay, extraction buffer (50 mM HEPES, 150 mM KCL, 1 mM EDTA, and 0.1% Triton X-100; pH 7.5), supplemented with 1 mM DL-Dithiothreitol (DTT) and protease inhibitor cocktail (Sigma-Aldrich, St. Louis, MO, USA), was used for protein extraction from plant materials. *N. benthamiana* leaves were frozen in liquid nitrogen and polished to a fine powder, add 1ml of the configured protein extract for every 0.5 g of leaves, vortex and mix well, incubate at 4 °C for 30 minutes to fully lyse the sample, centrifuge at 13000 rpm for 15min at 4 °C, take 80 ul of 5 x sample loading buffer, mixed well and then placed in a metal bath at 100 °C for 10min. When the extracted protein is used for Co-IP assay, the above method is used to sample for input in SDS-PAGE, and the remaining supernatant is incubated with target beads.

### Western blot

Proteins from the sample lysate were fractionated by sodium dodecyl sulfate polyarcylamide gel electrophoresis (SDS-PAGE). Sample in the gel were transferred to PVDF membrane using eBlot™ L1 (GenScript Corporation). Anti-HA (1:5, 000; #M20013; Abmart Inc., Shanghai, China), anti-flag (1:5, 000; #M20018; Abmart), antibodies were used to bind the protein with the corresponding tag. The total protein is displayed in this paper by ponceau staining.

### Luciferase complementation assay

The coding sequence of indicated genes was cloned into pCAMBIA1300-35S-HA-Nluc-RBS or pCAMBIA1300-35S-Cluc-RBS and then was transferred into *A. tumefaciens* strain GV3101. The constructs were co-expressed in *N. benthamiana* plants, OD_600_=0.5. The leaves were infiltrated with 1 mM luciferin (Biovision) at 2dpi, then the leaves were detected with a microplate reader (BioTek, Beijing, China). Two leaves were used for each test and three independent experiments were performed with same results.

### Yeast two-hybrid system

pGBKT7-Bait or pGADT7-Prey constructs used in S9 Fig and were transformed into the yeast strain AH109. Co-transformants were plated on double dropout supplements (SD-LW) and incubated at 28 degrees for 3days. After the yeast colonies on the Double Dropout Supplements plates grew, it indicated that the pGBKT7 and pGADT7 vectors were successfully co-transformed. Next, the yeast colonies were diluted with sterile water to OD600=0.1, followed by another 10-fold and 100-fold, respectively. Take 10 ul of yeast liquid droplets on the double dropout supplements (SD-LW) and triple dropout supplements (SD-LWH), and take pictures and records after incubating at 28 degrees for 3 days.

### Phylogenetic analysis

Multiple alignments of full-length amino acid sequences were aligned using MUSCLE. Phylogenetic analysis was performed using NB-ARC sequences with the fasttree program [58] or MEGA X program [59] by the maximum likelihood method with 100 bootstrap samples and parameters: poisson model, uniform rates and complete deletion. Alignments data and phylogenetic tree data could be found in S2 and S3 Files.

### Accession number

The primary accession codes for PN1a, PN1b, PN2a, PN2b, NRCX, NRCY that support the finding of this study were shown as NbD002668.1 (PN1a), NbD002669.1 (PN1b), NbD026701.1 (PN2a), NbD026704.1 (PN2b), NbD022578.1 (NRCY), NbD022579.1 (NRCX), respectively.

## Abbreviations

ETI: Effector-triggered immunity
HR: Hypersensitive response
CC: Coiled-coil
TIR: Toll/interleukin 1 receptor (TIR) domain
NRC:NB-ARC: Nucleotide Binding (NB)-ARC (APAF1, R gene products, and CED-4) domain
LRR: Leucine-rich repeat
NRC-H: NRC helper
ID: Integrated decoy
Sensor: An NLR that is responsible for identifying effectors
Helper: Downstream of sensor NLR-mediated immunity
Paired NLR: Two NLR genes, whose loci are very close to each other, work together to regulate immunity
SD Media: Synthetic Dropout Media.

## Supporting information

**S1 Fig. The pipeline of paired NLRs and NRC-Hs identification**.

NLR proteins were identified from plant proteome using NLRtracker. Sequences with NB-ARC annotated by NLRtracker were considered as NLRs. NLRs in “head-to-head” fashion with enclosing no more than 2 non-NLR genes are paired NLRs. NRC-Hs were identified by phylogenetic assay (S2 File). NLR clustered with functionally validated NRC helpers, NRC2/3/4, are NRC-Hs.

**S2 Fig. NLR identification materials and NLR evolutionary trees for different species**.

(A) Summary of plants species used in this study. Different branches of the species are represented by different colors corresponding to figure 1. (B) High similarity between NLR numbers identified in this study and those identified by Ngou et al.. Black line represents the linear trend, with dotted line representing the 95% confidence interval. Regression equation (with its R square value) and Person correlation value is shown at top right.

**S3 Fig. No correlation between %tail-to-tail NLRs and %head-to-head NLRs against %NRC-Hs in Solanaceae species**

(A-B) Scatter plot of %tail-to-tail NLRs and %head-to-head NLRs against %NRC-Hs. Black line represents the linear trend, with dotted line representing the 95% confidence interval. Regression equation (with its R square value) and Person correlation value is shown at top right.

**S4 Fig. Virus-induced silencing of *NRCX* impairs growth and promotes senescence in *N. benthamiana***.

(A) The silencing efficiency of indicated genes. TRV empty vector (TRV:*GUS*) was used as a negative control. (B) The morphology of 6-week-old *NRCX*-silenced *N. benthamiana* plants. Two-week-old *N. benthamiana* plants were infiltrated with *Agrobacterium* strains carrying tobacco rattle virus (TRV) VIGS constructs, and photographs were taken at 4 weeks after the agroinfiltration. Bar = 2 cm. (C,D) Accelerated senescence in *NRCX*-silenced plants. The fourth true leaf (counting from bottom to top) at 5-week-old plant was harvested (C) and the chlorophyll contents were calculated and shown in (D). Asterisks indicate statistically significant differences with. Three independent biological replicates were performed and indicated by different colored dots (** *P* < 0.01, *t* test; mean ± SD, n =12). Bar = 2 cm. (E) Virus content in TRV:*GUS* and TRV:*NRCX* was detected by qRT-PCR (** *P* < 0.01, *t* test; mean ± SEM, n =12).

**S5 Fig. No off-target effects in *nrcx* mutant**.

(A) Evolutionary tree of *NRCX* homologous genes in *N. benthamiana*. The confidence levels are marked in the node of branches. (B) Flanking sequence of potential CRISPR target. The *nrcx* mutant contains a 7 bp deletion in *NRCX* locus and not mutation could be found in other potential targets.

**S6 Fig. Silencing efficient of *NRCX* and *NRCY* in *N. benthamiana***.

(A) Gene fragments used for VIGS. The gene fragment used for VIGS are marked by light blue. (B) Silencing efficient of *NRCX* and *NRCY* in VIGS experiments (** *P* < 0.01, *t* test; mean ± SEM, n = 3) (C) Silencing efficient of *NRCX* and *NRCY* in RNAi experiments (* *P* < 0.05, ** *P* < 0.01, *t* test; mean ± SEM, n = 3).

**S7 Fig. The phylogenetic assay of NRCY**.

The maximum likelihood phylogenetic tree was generated in RAxML version 8.2.12 with JTT model using NB-ARC domain sequences of 2189 NLRs identified from tomato (Solyc-), N. benthamiana (NbD-), Coffee (Cc-), Kiwifruit (DTZ-), Sugar beet (EL-), Arabidopsis (AT-), Rice (Os-) (Sl Appendix, FileS3). NRCY-clade and NRCX-clade are marked by blue and orange, respectively. NRCY-clade are expanded and shown below. Domain architecture and motif sequences of NRCY clade members are shown at bottom right. Each motif was identified in MEME using NRCY-clade sequences.

**S8 Fig. NRCX negatively modulates NRC3-mediated cell death**.

(A) Hypersensitive cell death phenotypes. Infiltration site of leaves expressing NRC3^D480V^, NRC4^D478V^, Nsm/Sw5b and Avr3a/R3a were photographed at 5 days post infiltration and related cell death intensities were scored. (B) Calculated HR index of indicated leave sites. Three independent biological replicates were performed and indicated by different colored dot. Asterisks indicate statistically significant differences with *t* test (\*\**P* < 0.01). (C) The leave phenotype of each HR index. The scale representing cell death extent ranges from 0 (no cell death) to 8 (extensive cell death).

**S9 Fig. CC domains of NRCX and NRCY form heterodimer**.

(A) Schematic diagram of domain segmentation of NRCX and NRCY. (B) Verification of the interaction between CC domains of NRCX and NRCY by yeast two-hybrid (Y2H) assay. Ev/p53 is positive control for interaction. (C) Association between CC domains of NRCX and NRCY detected by luciferase complementation assays. FLS2-nLUC + cLUC-AGB1 was used as positive control.

**S1 File. List of primers in this article**.

**S2 File. NLR evolutionary tree files for 29 species**.

**S3 File. Evolutionary tree file of the gene *NRCY***.

## Declarations

### Ethics approval and consent to participate

Not applicable.

### Consent for publication

Not applicable.

### Availability of data and material

Not applicable.

### Competing interests

The authors declare that they have no competing interests.

### Funding

The work was supported by the National Natural Science Foundation of China (31625023 and 32072507) and the Fundamental Research Funds for the Central Universities (KYT202001 and JCQY201904).

### Authors’ contributions

DD, XD and GA conceived and designed the project, jointly performed data analysis and wrote the manuscript. XD, XZ, ZY, JL, CX, WP, TL, HZ, ZY, YZ, YL, ZW, ZK and YY performed the experiments. XD and GA analysed data. DD, XD and GA wrote and modified the manuscript. All authors read and approved the final manuscript.

## Acknowledgements

We appreciate Dr. Min Zhu and Prof. Xiaorong Tao at Nanjing Agricultural University for their gift of NRC3/4 auto-activated mutant, constructive suggestions and help. We appreciate Dr. Meixiang Zhang at Shaanxi Normal University for his help.

## Notes

### Competing Interest Statement

The authors have declared no competing interest.

